# Genomic erosion in the assessment of species’ extinction risk and recovery potential

**DOI:** 10.1101/2022.09.13.507768

**Authors:** Cock van Oosterhout, Samuel A. Speak, Thomas Birley, Lewis WG Hitchings, Chiara Bortoluzzi, Lawrence Percival-Alwyn, Lara Urban, Jim J. Groombridge, Gernot Segelbacher, Hernán E. Morales

## Abstract

Many species are undergoing rapid population declines and environmental deterioration, leading to genomic erosion, the loss of genetic diversity, accumulation of deleterious mutations, maladaptation, and introgression, that undermines fitness and long-term viability. Critically, this process continues even after demographic recovery due to a time-lagged genetic burden known as drift debt. Current conservation assessments, such as the IUCN Red List, focus on short-term extinction risk and do not capture the long-term consequences of genomic erosion. Likewise, the longer-term assessments of the IUCN Green Status may overestimate population recovery by failing to account for the lasting effects of genomic erosion. As genome sequencing becomes increasingly accessible, there is a growing opportunity to quantify genomic erosion and integrate it into conservation planning. Here, we use genomic simulations to illustrate how how different genomic metrics are sensitive to the drift debt. We test how ancestral effective population size (N_e_) and bottleneck history influence the tempo and severity of genomic erosion, and demonstrate how these dynamics shape genetic load and additive genetic variation, key indicators of long-term evolutionary potential. Finally, we present a proof-of-concept for a Genomic Green Status framework that aligns genomic metrics with conservation impact assessments, laying the foundation for genomics-informed strategies to support species recovery.

## Introduction

Conservation biology has long been characterized as a “mission-oriented crisis discipline,” in which management actions must be taken rapidly, often with limited data and resources (Soulé, 1985; McDonald-Madden *et al*., 2008; Wilson *et al*., 2011). Over the past decades, many species have been saved from extinction (Hoffmann, 2010; Bolam, 2021). However, as the biodiversity crisis accelerates, human-induced environmental changes are causing rapid population decline across taxa (Watson, 2019). Although demographic metrics such as census size and geographic range have guided most conservation policy to date, there is growing recognition that genetic factors critically influence species’ resilience, extinction risk, and capacity for recovery, particularly in the long-term (Frankham, 2005; Forester *et al*., 2022; Exposito-Alonso *et al*., 2022; Wilder *et al*., 2023; van Oosterhout, 2024; Shaw *et al*., 2025). This long-term perspective is of critical importance because current conservation and extinction-risk assessments, e.g. the IUCN Red List, focus on short-term dynamics over three years or ten generations (whichever is longest) (IUCN, 2004). Advances in genomic sequencing now allow us to analyse whole genomes to reconstruct recent changes in demography and evolutionary events, and to quantify genome-wide diversity and characterise functional and harmful genetic variation. These metrics can offer powerful insights into population viability and adaptive potential across long-term timescales (Soulé, 1987; Charlesworth, 2009; Lowe *et al*., 2017; Kardos, 2021; Moran, 2021; Forester *et al*., 2022; Willi, 2022). Despite this potential, genomic data remain peripheral in conservation and extinction-risk assessments, and the explicit protection of genetic diversity continues to lag behind species- and ecosystem-level priorities (Laikre, 2010; Willoughby, 2015; Hoban, 2023). This situation ultimately leaves a critical gap and fails to incorporate evolutionary processes into conservation planning (Hoffmann, 2015; Cook and Sgrò, 2019; Geue *et al*., 2025; Shaw *et al*., 2025).

Bridging this disconnect between short-term demographic focus and long-term genetic health calls for incorporating the concept of genomic erosion into conservation and extinction-risk assessments. We define genomic erosion as the progressive loss of genome-wide diversity over time, driven primarily by genetic drift and inbreeding in small or bottlenecked populations. This process undermines individual fitness, reduces long-term viability and adaptive potential, and ultimately elevates extinction risk. Importantly, genomic erosion often remains cryptic, because after demographic decline, the population’s new mutation-drift equilibrium is reached only slowly, leading to a prolonged “drift debt” (Gilroy *et al*., 2017; Dussex, Morales, *et al*., 2023; Pinto *et al*., 2024; Gargiulo *et al*., 2024; Liu *et al*., 2025).

We argue that understanding genomic erosion is essential for accurately assessing extinction risk and evaluating long-term recovery potential. We advocate for the stronger integration of genomic tools in conservation biology, emphasizing how recent advances in temporal genomics and simulation-based approaches can provide critical insights into extinction dynamics. These tools offer the opportunity to detect hidden genetic risks, forecast future trajectories, and improve the precision and effectiveness of conservation decisions in a time of accelerating global biodiversity loss.

### Genomic erosion

Genomic erosion is a pervasive – but frequently overlooked – consequence of the many threats faced by wild populations, such as overexploitation, invasive species, emerging infectious diseases, hybridisation, pollution, and habitat and environmental change. These threats fundamentally alter the strength and direction of evolutionary forces. Specifically, the gene pool of threatened populations may experience more genetic drift and novel selection pressures (Mooney and Cleland, 2001; Couvet, 2002; Fogell, 2021; Moran, 2021). Moreover, altered patterns of gene flow and recombination can result in the introgression of the genome by heterospecific DNA (Rhymer and Simberloff, 1996; Moran, 2021), whilst environmental pollution can increase the (germline) mutation rate (Somers *et al*., 2002; Keith, 2021). These genomic changes can reduce survival and reproduction, undermining overall population performance. The impacts of genomic erosion are manifested as: (1) a loss of genome-wide diversity reducing adaptive potential, (2) an elevated realised load (that is, the component of genetic load of harmful variants whose fitness effects are expressed), (3) a mismatch between genetic adaptations and the prevailing environmental conditions (i.e., maladaptation), and (4) the negative consequences of genetic introgression due to hybridisation.

Although rarely the sole cause of extinction, genomic erosion interacts with demographic decline, habitat degradation, and other stressors to drive populations into genetic Allee effects (Luque *et al*., 2016), mutational meltdown (Lynch *et al*., 1995), insufficient adaptive evolutionary potential (Forester *et al*., 2022), and an extinction vortex (Fagan and Holmes, 2006). Accordingly, genomic erosion often plays a critical role during the later stages of population decline, when the fate of a population or species is ultimately decided (Spielman *et al*., 2004). Moreover, the drift debt creates a time-lag in allele and genotype frequency changes, imposing a hidden genetic burden that may only become apparent several generations after the initial disturbance (Jackson *et al*., 2022; Pinto *et al*., 2024; Liu *et al*., 2025). This time-lag is particularly problematic for conservation assessments because populations may initially appear genetically healthy right after population decline. Crucially, previously bottlenecked populations that show partial demographic recovery are often downlisted in the IUCN Red List. Yet, these populations may continue to lose genetic diversity due to drift debt, thereby increasing their extinction risk (Jackson *et al*., 2022; Fontsere *et al*., 2024). Additionally, some conservation actions can exacerbate genomic erosion, for example by relaxing selection pressures through supplementary feeding in the wild or captive breeding in zoos (Araki *et al*., 2007; Frankham, 2008; Robinson *et al*., 2023). Thus, even after immediate threats are mitigated, genomic erosion can persist as a long-term constraint on recovery and viability, potentially causing the Red List assessment to underestimate extinction risk.

To illustrate the impact of drift debt on genomic erosion, we performed forward-in-time computer simulations under a range of bottleneck and population recovery scenarios (Fig. 1A). The timing and magnitude of responses varied across genetic metrics and bottleneck intensities. Among diversity statistics, nucleotide diversity (*π*) responded slowly, exhibiting a pronounced drift debt as it continued to decline for several generations after demographic recovery (Fig. 1B). The number of segregating sites (*S*) responded more rapidly, reflecting the swift loss of rare alleles, and stabilized shortly after the recovery phase (Fig. 1C). As a consequence, Tajima’s *D* became strongly positive during the early recovery period, indicating a skewed allele frequency spectrum (Fig. 1D). Genetic load metrics also exhibited distinct temporal dynamics under the same demographic dynamics. Realized load surged rapidly following the crash, especially during the first 10–20 generations when populations are most vulnerable to inbreeding depression and extinction (Fig. 1E). In contrast, masked load declined more gradually during the bottleneck due to purging and conversion into realized load (Fig. 1E). Together, these results reveal that different metrics capture distinct phases of genomic erosion, with genetic diversity loss and elevated realized load persisting long after demographic recovery. This highlights the value of using multiple genomic indicators, and ideally temporal genomic data, to capture both immediate threats to viability and long-term adaptive capacity.

**Figure 1.**
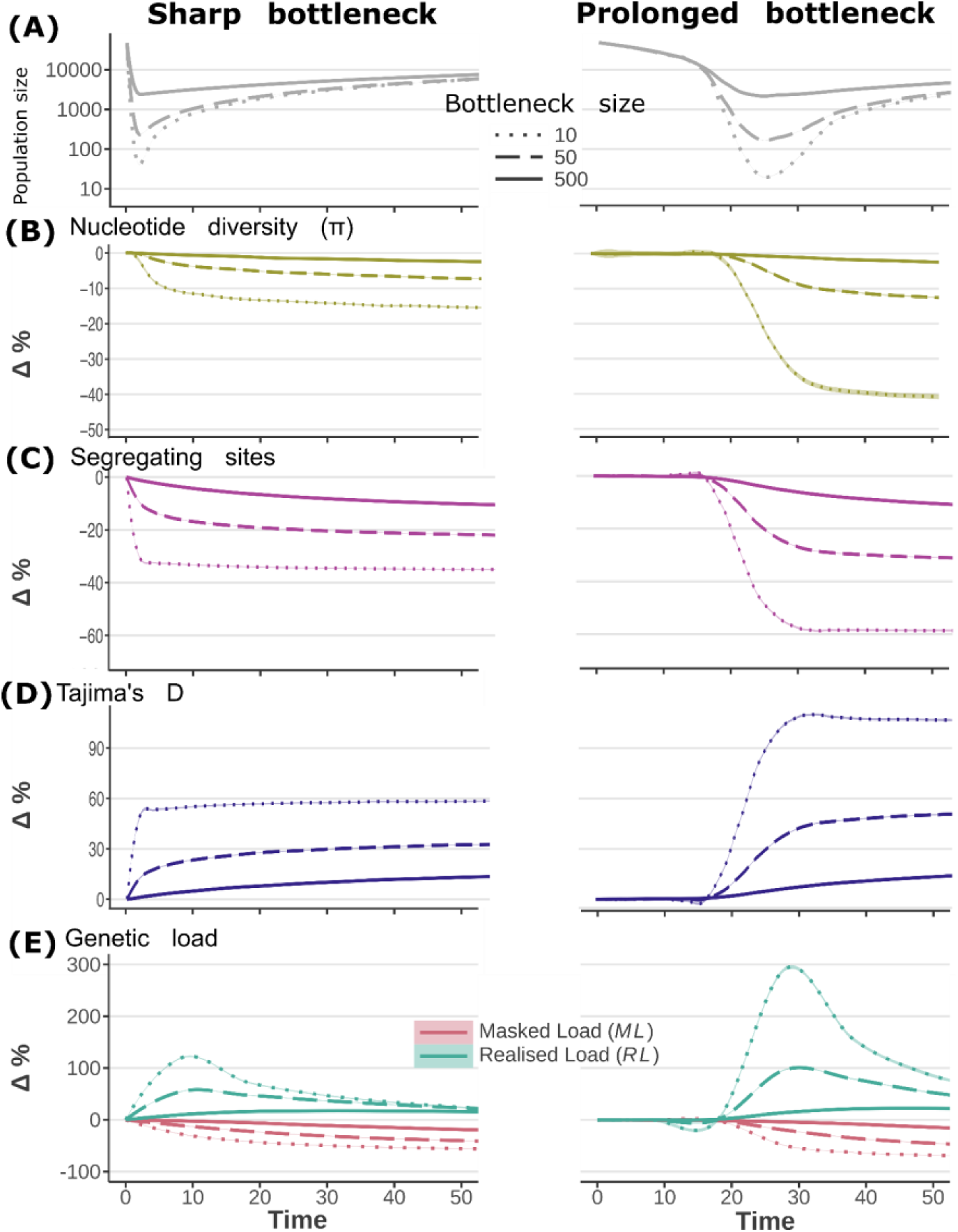
Genomic erosion metrics during population decline. Populations with an ancestral size of 10,000 breeding individuals were simulated in SLiM across contrasting bottleneck scenarios. The left panels depict fast and short bottlenecks (rapid decline over 1 generation and recovery after 2 generations), whereas the right panels show more gradual and prolonged bottlenecks (decline over 20 generations, recovery after 10 generations). (A) Demographic trajectories for three bottleneck intensities (Ne = 10, 50, 500). Panels (B–E) illustrate the temporal dynamics of five genomic metrics: (B) Nucleotide diversity (π), (C) Segregating sites (S), (D) Tajima’s D, and (E) Genetic load components (realised load RL and masked load ML). LOESS smoothing was applied with a span of 0.05 for metrics sampled every generation, and 0.1 for genetic load metrics (sampled every 10 generations) to reduce overfitting. These results highlight the contrasting temporal responses of genomic indicators to demographic change and the drift debt, revealing complementary insights into both immediate and delayed consequences of genomic erosion.

### The importance of the effective population size (N_e_)

Conservation assessments have traditionally been focused on census population size (N) and geographic range, yet effective population size (Nₑ) ultimately governs the balance between mutation, drift and selection. As such, Nₑ shapes both the retention of adaptive variation and the accumulation of deleterious alleles (Frankham, 2021; Laikre, 2021; Waples, 2025). Recently, effective population size (Ne) has been proposed as a genetic indicator for inclusion in the global biodiversity framework of the Convention on Biological Diversity (CBD), and as one of the genetic Essential Biodiversity Variables (EBVs) by the Group on Earth Observations Biodiversity Observation Network (GEO BON). However, many challenges remain in estimating N_e_ (Ryman *et al*., 2019; Fedorca *et al*., 2024). Confusingly, the N_e_ is an umbrella term that reflects the impact of genetic drift and inbreeding on different population genetic statistics (Waples, 2025). In practice this means that populations can have multiple, sometimes markedly different, N_e_ values depending on how this is estimated, such as the inbreeding N_e_, variance N_e_, and coalescence N_e_, which have been comprehensively reviewed in (Waples, 2025). This variety in N_e_ estimators can lead to misguided interpretations for conservation. Moreover, N_e_ estimators have different temporal resolutions from the ancestral N_e_ (thousands of generations ago), the recent N_e_ (one to hundreds of generations ago), and the current N_e_ (Nadachowska-Brzyska *et al*., 2022).

#### Ancestral N_e_

When N_e_ estimation is based on nucleotide diversity or the coalescence of alleles, it is largely shaped by the genetic effective size of the ancestral population many thousands of generations in the past. This ancestral N_e_ relates to the equilibrium between the input of genetic variation by mutations and loss of this diversity due to genetic drift (*Θ*=4N_e_μ). Software such as PSMC, MSMC, and Bayesian Skyline Plots (BSP) are commonly used to reconstruct historical demography based on the coalescence of alleles. However, high recombination rates (relative to mutation rates) can make N_e_ inference unreliable (Bortoluzzi *et al*., 2023). Furthermore, recent population size declines reduce genetic diversity, but the coalescence of alleles and loss of nucleotide diversity are slow processes that are markedly affected by the drift debt (Fig 1). Consequently, knowledge about ancestral N_e_ alone is of limited relevance for present-day extinction risk assessment of threatened species without proper context (see below).

#### Recent N_e_

Linkage-disequilibrium (LD) N**_e_** estimates respond more quickly to changes in population size as they reflect the evolutionary balance between recombination and inbreeding that is shaped by recent changes in demography over the past few hundred generations. Recombination reduces LD, whereas inbreeding (and hence, small N_e_) increases LD. Software such as GONE and SNeP can be used to infer this linkage-based estimate of N_e_, which can capture recent demographic events such as bottlenecks and founder events. Given that issues relating to inbreeding depression and the spike in realised load play out across this relatively recent timescale (Fig. 1), this makes the recent N_e_ more directly relevant to conservation assessments.

#### Contemporary N_e_

The most immediate estimate of N_e_ is based on loss in heterozygosity across one (or multiple) generations. For conservation genetic purposes, the contemporary N_e_ estimate is particularly useful because it reflects the status of diversity loss in the current population relative to a previous sample. Unfortunately, it requires temporal genomic samples from two (or more) generations, which has thus far has limited its application. Moreover, admixture between previously isolated populations or distinct evolutionary significant units (ESUs) can artificially inflate contemporary N_e_ estimates, which emphasizes the importance of assessing the loss of diversity within ESUs (Geue *et al*., 2025).

Increasingly, conservation genetic studies report the N_e_/N_c_ ratio (Waples, 2024). This ratio is inflated in many threatened species, especially in recently bottlenecked populations (Wilder *et al*., 2023; Wang *et al*., 2025). However, it is critically important to know how the N_e_ was estimated. In species that experienced a gradual decline in population size, the N_e_/N_c_ ratio based on ancestral N_e_ estimators is likely to be significantly inflated due to the drift debt. Afterall, nucleotide diversity and the coalescence of alleles changes only slowly during population size decline. Hence, the ancestral N_e_ lacks behind the census population size (N_c_), inflating the N_e_/N_c_ ratio (Wilder *et al*., 2023; Waples, 2024; Wang *et al*., 2025). However, using a recent or current N_e_ estimate, the N_e_/N_c_ ratio is likely to be much less inflated by the drift debt.

### Genetic load of deleterious mutations

Populations with large effective population sizes (Nₑ) are expected to accumulate a substantial genetic load of partially recessive deleterious mutations at low frequency. These variants comprise the masked load, as their fitness effects remain largely hidden from selection while present in heterozygous form (Bertorelle, 2022). The masked load does not reduce mean fitness. However, when population size declines and inbreeding increases, homozygosity rises, converting masked load into realized load (García-Dorado, 2012; Hedrick and Garcia-Dorado, 2016; Dussex, Morales, *et al*., 2023), which exposes deleterious effects and leads to inbreeding depression (Hedrick and Garcia-Dorado, 2016; Smeds and Ellegren, 2022).

Accurate extinction risk assessment therefore requires reconstructing the demographic trajectory of Nₑ over time (Fig. 2A). Populations with large ancestral Nₑ accumulate more deleterious alleles as masked load and are at greater risk of severe inbreeding depression following demographic collapse (Grossen *et al*., 2020; Bertorelle, 2022; Kleinman-Ruiz, 2022; Femerling *et al*., 2023; Dussex, Morales, *et al*., 2023). Forward-in-time simulations illustrate that ancestrally large populations harbor more nucleotide diversity (Fig. 2B) and higher masked load (Fig. 2E). After undergoing an equivalent severe bottleneck (to Nₑ = 10), these populations convert substantially more masked load into realized load (Fig. 2F), resulting in markedly elevated extinction rates (Fig. 2C) compared to populations with smaller ancestral Nₑ. Thus, large ancestral Nₑ can be a red flag for declining species, including zoo populations derived from a small number of founders.

**Figure 2.**
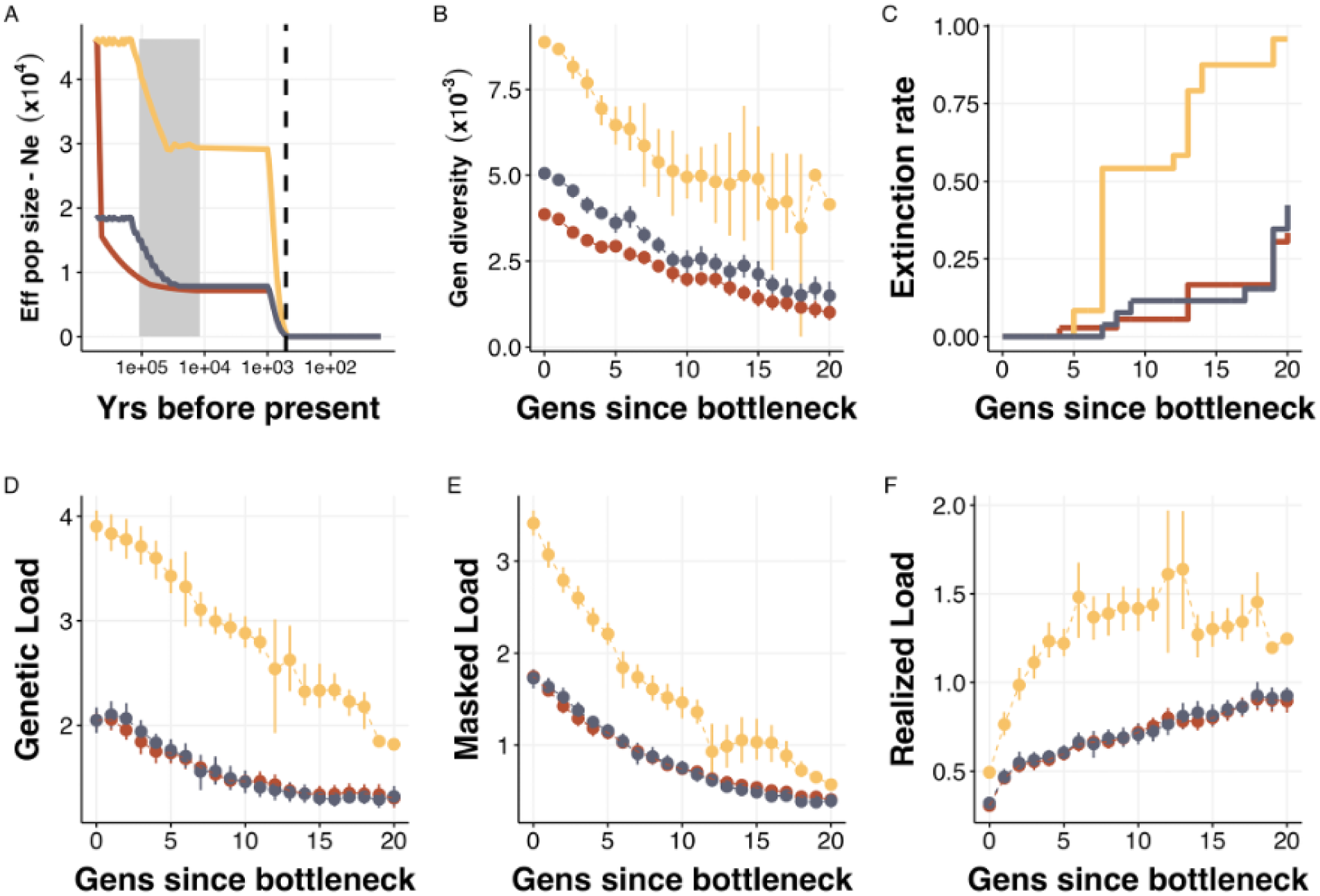
The effects of ancestral population size on genetic load dynamics. (A) Populations with distinctly different ancestral demographic trajectories experienced a severe population bottleneck (Ne=10). Grey shading represents the Last Glacial Period 110–12 thousand years ago. Dotted line represents the beginning of the Anthropocene in the year 1610. (B) The ancestrally large populations (yellow) show the highest nucleotide variation, but panel (C) shows that such populations also have the highest extinction rate after a bottleneck. (D) This is because the genetic load from unconditional deleterious mutation is highest in the ancestrally large populations (yellow). (E) Historically, when the population was still large, the genetic load was not expressed, and this part of the genetic load is known as the masked load. (F) However, population size decline results in inbreeding, during which the masked load is converted into a realised load.

In addition to simulations, genomic data can be used to reconstruct historical demography and estimate genetic load across the genome. Advances in genome annotation and functional prediction (e.g., tools like CADD and GERP) allow estimation of deleterious variant burden (Kircher, 2014; Bertorelle, 2022; Speak *et al*., 2024). Predictions validated in model species can be transferred to threatened taxa (Fontsere *et al*., 2024; Speak *et al*., 2024; Wang *et al*., 2025), and emerging deep learning models further improve prediction accuracy (Frazer *et al*., 2021). Comparative genomic analyses confirm that genetic load scales with ancestral Nₑ, and that populations with recent declines often suffer from elevated realized load (Wang *et al*., 2025). Temporal genomic datasets are especially valuable, enabling direct observation of masked-to-realized load conversion and providing a dynamic framework for assessing the impact of genomic erosion over time (Van Der Valk *et al*., 2019; Dussex, 2021; Dussex, Kurland, *et al*., 2023; Femerling *et al*., 2023; Bortoluzzi *et al*., 2024; Fontsere *et al*., 2024; Cavill *et al*., 2024).

### Maladaptation and adaptive potential

Polymorphisms at quantitative trait loci (QTL) can be either deleterious or beneficial depending on the genetic background and environmental conditions (Charlesworth, 2013a, 2013b; Kardos, 2021). Most outbred populations are adapted to their environmental optimum as additive genetic variation at QTL is maintained by stabilising selection acting on the trait (Charlesworth, 2013a). Thus, genomic erosion could lead to maladaptation by removing additive genetic variation, and the outcome of this process depends on the ancestral N_e_ (Fig. 3). Perhaps counterintuitively, populations with a large ancestral N_e_ have on average a lower fitness from traits under stabilising selection (Fig. 3) because larger populations are closer to the trait optimum so any new mutation will be (on average) more deleterious (Charlesworth, 2013a). However, the amount of additive genetic variation segregating in the ancestral population also underpins their adaptive evolutionary response during environmental change. Hence, populations with large ancestral N_e_ are better able to respond to environmental change, and theoretically, they are expected to have a lower extinction risk than small populations (Fig. 3). On the other hand, as previously shown, during population decline ancestrally large populations are more prone to inbreeding depression due to a higher masked genetic load (Fig 3). Thus, high ancestral Nₑ is a double-edged sword: it promotes historical diversity but allows deleterious variants to persist at low frequency, only to manifest as inbreeding depression and elevated extinction risk under collapse. Moreover, maladaptation may also arise from gene–environment mismatches, particularly under climate change, where formerly adaptive traits may become deleterious in novel environmental conditions. This highlights the importance of assessing the multifarious threats of genomic erosion using computer models that incorporate the effects of the demographic history, different types of genetic variation, selection regimes and realistic rates of environmental change (Forester *et al*., 2022; Jackson *et al*., 2022; Robinson *et al*., 2022; Nigenda-Morales *et al*., 2023; Pinto *et al*., 2024).

**Figure 3.**
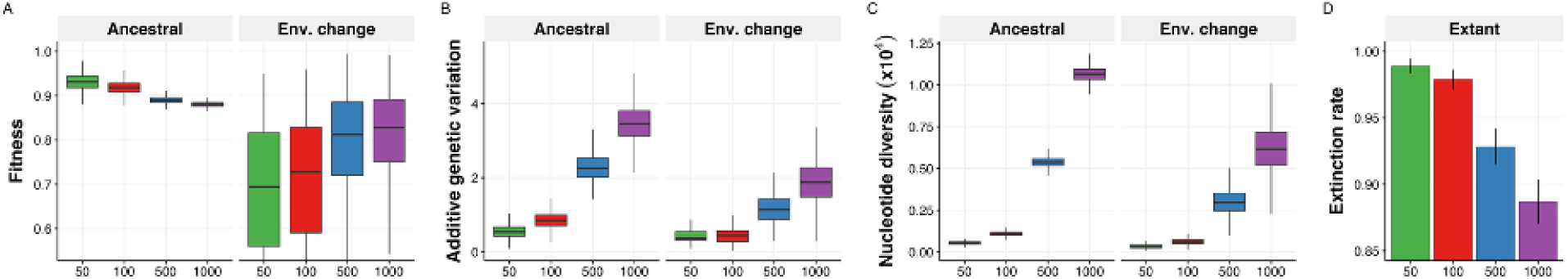
The effects of ancestral population size on adaptation. Computer simulations in SLiM of populations with different ancestral effective population sizes (Ne=50, Ne=100, Ne=500, Ne=1000) with a trait adapted to an environmental optimum. The populations experience a severe population bottleneck (*Ne*=10) that reduces genetic diversity, and five generations after experience an environmental change that shifts the optimum of the trait value. Here we show the distribution of values of across five generations before the population bottleneck (ancestral population) and five generation following the optimum shift. The larger ancestral populations have a slightly lower fitness **(A)** because they possess more additive genetic variance (V_A_) **(B)**, mirrored by a higher amount of genome-wide genetic diversity **(C)**. This results in more phenotypic variation around the optimum and it constitutes a genetic load of conditionally deleterious mutations. However, this larger additive variation becomes beneficial when the environmental optimum changes as it underpins the adaptive response to selection. Consequently, larger ancestral populations have a higher fitness after the optimum shift and a lower extinction rate **(D).** V_A_ is positively correlated to neutral genetic diversity, highlighting the value of high genetic diversity to preserve adaptive potential. For simplicity, we simulated a single additive polygenic trait without environmental variance to illustrate the reduction of V_A_. Parameters such as dominance, epistasis and the genetic architecture of the trait might temporarily increase V_A_ after a bottleneck (Goodnight, 1988; Willis and Orr, 1993; Barton and Turelli, 2004). However, over time, genetic drift is expected to lead to a reduce adaptive response under most conditions.

Similarly, the loss of immunogenetic diversity constitutes a key component of genomic erosion. The major histocompatibility complex (MHC) and toll-like receptors (TLRs) are among the most studied immune loci, and are typically subject to balancing selection that maintains high allelic diversity within populations (van Oosterhout, 2009; Spurgin and Richardson, 2010; Gilroy *et al*., 2017). However, bottlenecked populations can lose immunogenetic diversity, which makes them more susceptible to disease outbreaks (Grueber *et al*., 2012; Morris *et al*., 2015; Dalton *et al*., 2016; Fogell, 2021; Silver *et al*., 2025). Erosion of immunogenetic diversity does not necessarily proceed at a similar rate as neutral diversity (Lighten *et al*., 2017; Gilroy *et al*., 2017), which highlights the need to monitor functional loci alongside genome-wide markers. Maintaining immunogenetic diversity is critical for managing disease risk in small populations, guiding translocations, and identifying targets for gene editing aimed at restoring functional variation (Morris *et al*., 2015; Silver *et al*., 2025; van Oosterhout *et al*., 2025).

### Genomics-informed computer modelling

To assess the long-term risks posed by genomic erosion, conservation efforts must integrate genomic data analysis with computer modelling (Kardos, 2021; Kyriazis *et al*., 2023; Mathur *et al*., 2023; Dussex, Morales, *et al*., 2023). For decades, conservation scientists have relied on population viability analyses (PVA) to assess threats and inform conservation actions (Lacy, 2019). Although traditional PVA models can incorporate genetic data to assess the effects of inbreeding on population viability, they were not designed to capture genome-wide patterns of erosion. The value of evolutionary theory, computer modelling, and genomics is increasingly recognized in conservation, with the latter two fields advancing especially rapidly (Frankham *et al*., 2019; Funk *et al*., 2019; Hohenlohe *et al*., 2021; Segelbacher, 2022; Willi, 2022; Shaw *et al*., 2025). A new generation of evolutionary genomics models enables the construction of complex, genome-scale simulations that integrate demographic, ecological, and evolutionary dynamics within a unified framework (Guillaume and Rougemont, 2006; Haller and Messer, 2019, 2023; Terasaki Hart *et al*., 2021) (Fig. 1).

Genomic data provide the foundation for these models. Demographic inferences, mutation rates (e.g., from parent-offspring trios (Bergeron *et al*., 2023)), and recombination landscapes (Peñalba and Wolf, 2020) can be used to parameterise realistic genome architectures. These models can also incorporate species-specific traits such as reproductive strategy, dispersal, and longevity, and be made spatially explicit to assess the impact of metapopulation dynamics or habitat fragmentation (Pinto *et al*., 2024). Crucially, simulated outcomes can be compared directly to empirical genomic datasets for validation, enabling predictions under different environmental and management scenarios.

Forward-in-time simulations are increasingly applied to forecast the impacts of climate change, land-use change, loss of connectivity, adaptive potential, and genetic rescue (Matz *et al*., 2018; Brauer and Beheregaray, 2020; Dussex, 2021; Hansson *et al*., 2021; Kyriazis *et al*., 2021; Stoffel *et al*., 2021a; Jackson *et al*., 2022; Magliolo, 2022; Beichman, 2023; Femerling *et al*., 2023; Kyriazis, 2023; Kyriazis *et al*., 2023; Dussex, Morales, *et al*., 2023; Al Hikmani *et al*., 2024; Pinto *et al*., 2024; Cavill *et al*., 2024; Liu *et al*., 2025). Yet challenges remain, accurate model parameterisation requires genomic and ecological data that are still lacking for many species, and comparative frameworks are only now emerging. Increasing availability of temporal genomic datasets opens a powerful opportunity to validate forward-in-time simulations by directly comparing simulated genomic trajectories to observed changes over time. Such validation allows researchers to assess whether simulations accurately capture the pace and magnitude of genetic erosion following demographic collapse or recovery, including shifts in heterozygosity, allele frequency spectra, or realized load. Incorporating fitness data alongside genomic metrics further strengthens this framework, enabling the evaluation of how well predicted genetic load or inbreeding depression translates into fitness declines in real populations. As the volume and resolution of genomic time series increase, so too does the ability to calibrate models not only on past dynamics but to project genetic outcomes under alternative management actions or future environmental change. This approach transforms simulations from abstract scenarios into empirical, testable tools for forecasting extinction risk, recovery potential, and the long-term consequences of conservation interventions.

### Genetic risks and conservation assessments

The IUCN Red List remains the central global framework for assessing extinction risk, but its focus on short-term, immediate threats limits its ability to detect slower, longer-term processes of genomic erosion. There is mixed evidence as to whether Red List categories consistently reflect underlying levels of genetic diversity. Some studies show that threatened species tend to have lower genetic diversity than non-threatened species, but this pattern is not universal and appears to vary depending on the taxa, genetic markers and metrics used (Willoughby, 2015; Brüniche-Olsen *et al*., 2021; Canteri, 2021; Schmidt *et al*., 2022; Jeon *et al*., 2024; McLaughlin *et al*., 2025; Wang *et al*., 2025). Recent efforts have called for harmonized workflows and core genomic metrics, highlighting the importance of consistency in data generation and the selection of biologically meaningful, conservation-relevant indicators (Buzan *et al*., 2024; Jeon *et al*., 2024; McLaughlin *et al*., 2025).

Inconsistencies have sparked debate over whether the Red List can effectively protect intraspecific genetic diversity. Conversely, others have questioned whether given this poor association, genetic data can be used to assess extinction risk. We argue that both sets of data, ecological and demographic data collated in the Red List, and genetic or genomic data, are complementary, and that cover each other’s blind spots (van Oosterhout, 2024). Neither the Red List nor the Green Status of Species assess the impacts of genomic erosion. We stress that including genomic data in the Red List is critical because the loss of adaptive potential in combination with rapid environmental change poses unprecedented threats to wildlife. We therefore must assess the long-term viability of populations and species against the backdrop of environmental change, which requires analyses of genomic data, forward-in-time computer simulations, and Deep Learning models to decipher signals associated with elevated risk of extinction (van Oosterhout, 2024).

### Genomic Green Status

The International Union for Conservation of Nature (IUCN) recently developed the Green Status of Species, which is a framework for measuring species recovery and conservation impact (Akçakaya, 2018; Grace *et al*., 2021). The assessment calculates a Green Score that quantifies the viability, functionality and representation of a species, and this metric ranges between 0% (extinct) to 100% (fully recovered). The Green Scores are measured at four different timepoints (ancestral, current, next 10 years, and next 100 years), and differences in the Green Scores between those timepoints are used to estimate four conservation impact metrics. *Conservation Legacy* estimates the impact of past conservation on the population or species, comparing it to a counterfactual scenario without any conservation actions. *Conservation Dependence* assesses how much worse the species is likely to be after 10 years without any conservation. Conversely, *Conservation Gain* measures the potential improvement of the species after 10 years with conservation actions. Finally, *Recovery Potential* aims to assess the long-term improvement that could be accomplished during 100 years with continued conservation.

Given its longer timeframe, the Green Status of species could incorporate the dynamics of genomic erosion, consistent with the time-lag and long-term effects of the drift debt. Importantly, it also offers a clear conceptual framework for the integration of genomic data because its four conservation impact metrics are dimensionless units with scalar property. In other words, the percentages of metrics relating to different aspects of the species (e.g., genome-wide diversity, genetic load, individual fitness, etc.) can be directly compared across time and species. Importantly, the conservation metrics are proportional statistics that measure the change expected under a hypothetical scenario relative to the status of the species at present. Therefore, genomic indicators can be directly aligned with the Green Status metrics of *Recovery Potential* and *Conservation Gain*, offering a means to quantify the genomic dimension of long-term species extinction risk and recovery potential.

To demonstrate how genomic data can be integrated into the IUCN Green Status framework, we developed a simulation-based framework to adapt the current implementation focusing on demographic change to instead quantify genomic recovery using indicators of genetic load and genome-wide diversity. We use the pink pigeon (*Nesoenas mayeri*) as an example of a species that underwent a severe bottleneck (N∼12 during 1990’s) followed by a demographic recovery through intensive conservation (currently N∼488 adult birds). However, due to the drift debt, its long-term survival is threatened by genomic erosion (Jackson *et al*., 2022). The simulation model captures the species’ historical demography and recent management interventions, including genetic and demographic rescue from a captive population founded by 12 individuals in the 1970s, as implemented in Jackson et al. (2022).

We modeled four scenarios: (1) no conservation (counterfactual), (2) demographic rescue only, (3) genetic rescue only, and (4) combined demographic + genetic rescue. For each, we tracked realized genetic load, nucleotide diversity and extinction rates across time (Fig. 4). We adapted the Green Scores to calculate Species Recovery Scores (SRS) based on realized load and genome-wide diversity. For the realized load, the current SRS is 88.3%, indicating an elevated and sustained effect of harmful variation compared to the ancestral population, but when compared to the counterfactual scenario, the *Conservation Legacy* shows that conservation interventions likely saved the species from extinction. *Conservation Dependence* is high (52.6%), showing that ongoing genetic supplementation remains crucial. *Recovery Potential* is also substantial (23.5%), as continued management is expected to reduce realized load below ancestral levels via purging. In contrast, the Green Status for nucleotide diversity shows a lower SRS (25.4%), with modest Conservation Dependency (6.0%) and negative Recovery Potential (–8.7%), indicating that diversity loss is largely irreversible under current conservation scenarios.

**Figure 4.**
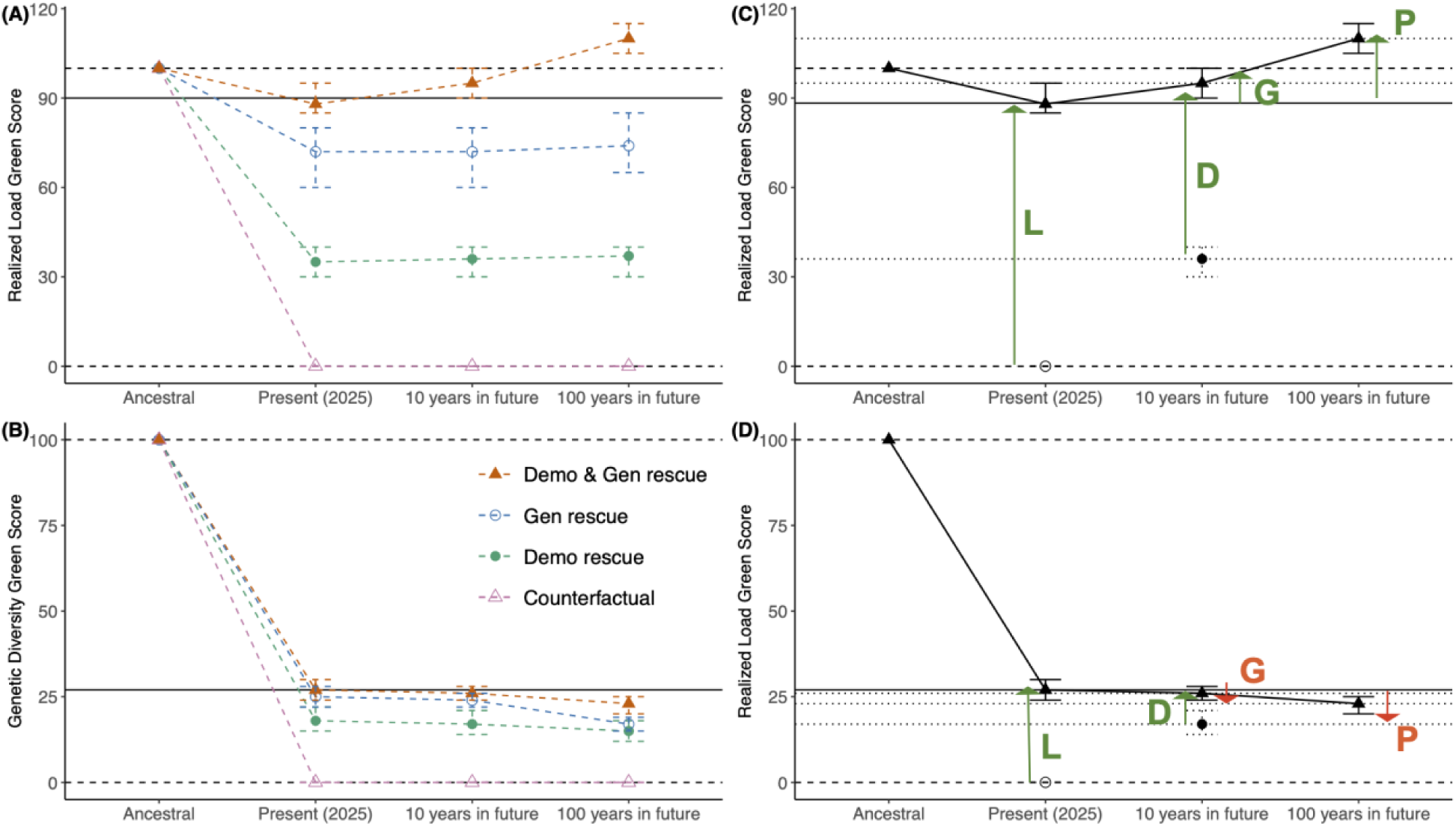
Genomic Green Scores for realized load and genetic diversity across four simulated conservation scenarios. The left panels show the change in (A) realized load and (B) genome-wide diversity over time for four conservation scenarios: (1) no conservation (counterfactual), (2) demographic rescue only, (3) genetic rescue only, and (4) combined demographic + genetic rescue. The right panels show the Genomic Green Scores for (C) realized load and (D) genome-wide diversity for the best-performing scenario of combined demographic + genetic rescue. Arrows show the change in species recovery scores (SRS) across time for Conservation Legacy (L), Conservation Dependence (D), Conservation Gain (G), and Recovery Potential (P). The change relative to the ancestral shows negative effects, where the realized load show a predicted decrease in fitness due to the accumulation of harmful variation, and the genome-wide diversity shows a decrease in genetic diversity as a proxy for adaptive potential.

By comparison, according to its Green Status assessment from 2021 (https://www.iucnredlist.org/species/22690392/179390191), the Species Recovery Score (SRS) for the Pink Pigeon is 17% (categorized as “Critically Depleted”), primarily reflecting extensive forest loss across its original range. It demonstrates a *High Conservation Legacy* between 14 and 17%—meaning that without prior conservation efforts, the species would very likely be extinct today. Its *Conservation Dependence* is *Low* between 5 and 8%, which means that if conservation efforts ceased, ecological functionality would deteriorate over a decade. Projected *Conservation Gain* over ten years is also *Low* and between 5 and 17%, indicating that the species has limited potential recovery. However, its long-term *Recovery Potential* over a 100-year timeframe is significantly higher, scoring between 15 and 36%, implying that restoration of sufficient habitat could allow ecological functionality in many areas. Taken together, the ecological and genomics Green Status assessment are complementary. By evaluating the genetic heath of the species, we believe the genomics Green Status assessment is also critical, as it identifies the urgent need for genetic rescue.

### The future of genomics-informed conservation

The study of genomic erosion is a rapidly advancing field, yet it faces several key challenges. Recent genomic analyses have shed light on how population declines and recoveries affect the balance between purging and accumulation of deleterious mutations, potentially shaping long-term fitness and viability in small populations (Grossen *et al*., 2020; Dussex, 2021; Humble, 2022; Kleinman-Ruiz, 2022; Riaño, 2022; Smeds and Ellegren, 2022; Dussex, Kurland, *et al*., 2023; Femerling *et al*., 2023; Kyriazis, 2023; Mathur *et al*., 2023; Fontsere *et al*., 2024). However, estimating additive genetic variation and directly linking genetic load with fitness effects remains difficult, especially in wild populations. Long-term monitoring programs, particularly those that incorporate fitness and genomic data across generations, offer one of the most promising avenues for quantifying adaptive potential and predicting extinction risk (Harrisson *et al*., 2019; Villemereuil, 2019; Fogell, 2021; Stoffel *et al*., 2021b; Bonnet, 2022; Jackson *et al*., 2022; Smeds and Ellegren, 2022; Kardos *et al*., 2023; Hewett *et al*., 2024; Morales, Norris, *et al*., 2024; Morales, Van Oosterhout, *et al*., 2024; Morales, Groombridge, *et al*., 2024).

Genomics-informed management is poised to become central to the conservation of both wild and captive populations. Zoo populations, often founded by very few individuals and exposed to relaxed selection in artificial environments, are especially vulnerable to genomic erosion. Genomic tools can help avoid unintended hybridization, limit the fixation of deleterious mutations, and detect adaptation to captivity. By monitoring allele frequency changes and minimizing artificial selection, genomic screening can reduce maladaptation and improve the success of reintroductions into the wild (Schulte-Hostedde and Mastromonaco, 2015). Similarly, in genetic rescue programs, targeted genome-wide screening enables the identification of optimal individuals that maximize diversity while minimizing the introduction of harmful mutations (Ralls *et al*., 2020; Kyriazis *et al*., 2021; Mathur *et al*., 2023; Speak *et al*., 2024). Several existing initiatives, such as the integration of genomic data into the Zoological Information Management System (ZIMS), and the development of large-scale biobanks, are already laying the groundwork for the systematic incorporation of genomic data into conservation practice (Schwartz *et al*., 2017; Pérez-Espona and CryoArks Consortium, 2021; Mooney *et al*., 2023).

To translate genomic insights into conservation practice, efforts must focus on developing standardized frameworks for calculating and reporting genomic erosion across taxa, including historical baselines derived from museum samples (Díez-del-Molino *et al*., 2017; Buzan *et al*., 2024). Importantly, common metrics that capture genetic load, diversity, and adaptive potential, should be defined to enable meaningful cross-species comparisons to guide conservation priorities (Jeon *et al*., 2024; Wang *et al*., 2025). The technical complexity of genomic analyses also requires harmonized pipelines and collaborative infrastructure, ensuring that conservation biologists, genomicists, bioinformaticians, and modelers can work together effectively.

Ultimately, a genomics-informed approach will allow conservation science to move from descriptive diagnostics to predictive frameworks. This shift will improve our ability to forecast extinction risk, measure conservation impact, and design recovery plans that secure the genetic health and evolutionary potential of species for generations to come.

## Methods

### Population bottleneck simulations

Simulations were performed in SLiM4 (Haller and Messer, 2023) with a non-Wright-Fisher model adapted to non-overlapping generations and random mating for simplicity. We simulated a genomic region modelled after chromosome 23 of the collared flycatcher genome (12.3 Mb) (Kawakami *et al*., 2014), incorporating realistic exon, intron, and intergenic region positions, as well as an underlying recombination map, thereby accurately representing linkage dynamics. We also simulated an exome architecture of 5 autosomes, each containing 1500 genes of 1500 bp with recombination rates of 1e-8 within genes and 1e-3 between genes. A global mutation rate of 1.5e-8 was used. Both neutral and deleterious mutations were simulated in ratios of 5:1 for introns, 1:2.31 for exons, and 1:0 for non-coding regions. Deleterious selection coefficients (s) were taken from a gamma distribution (mean=-0.05 and shape=0.5) with a tail of 5% of lethal mutation and negative relationship between s and dominance coefficients (h), following (Kardos *et al*., 2021)

The population size was controlled by limiting the number of breeding individuals each generation, with each breeding pair producing 12 offspring. Populations all had an ancestral size limited to 10,000 breeding individuals. We explored different bottleneck decline speeds (1 and 20 generations), bottleneck durations (2 and 10 generations), and bottleneck sizes (10, 20, 50, 100, and 500 breeding individuals) with each combination of parameters being run for 100 replicates.

### Genetic load simulations

We used the same modelling approach as in (Dussex, Morales, *et al*., 2023). Briefly, simulations were performed in SLiM3 (Haller and Messer, 2019) with a non-Wright-Fisher model adapted to non-overlapping generations and random mating for simplicity. The model simulated an exome of 3000 genes of 3.4Kb each with a recombination rate r=1e-4 (no recombination within genes), and a per base mutation rate m=1.4e-8. Deleterious selection coefficients (s) were taken from a gamma distribution (mean=-0.05 and shape=0.5) with a tail of 5% of lethal mutation and negative relationship between s and dominance coefficients (h), following Kardos et al., 2021. We ran 100 replicates per scenario.

### Additive genetic variation simulations

We used the same modelling approach as in (Femerling *et al*., 2023). Briefly, simulations were performed in SLiM3 (Haller and Messer, 2019) with a non-Wright-Fisher model adapted to non-overlapping generations and random mating for simplicity. The model simulated an exome of 3000 genes of 3.4Kb each with a recombination rate r=1e^-4^ (no recombination within genes), and a per base mutation rate m=1e^-7^. Fitness was determined based on the additive effect of genotype values (z) on a polygenic trait tracking an environmental optimum (opt) following (Falconer and Mackay, 1996). Genotype values (z) were drawn from a uniform distribution from −0.5 to 0.5 and had a fixed additive effect (h=0.5). The phenotype (P) of an individual was the sum of all homozygous and heterozygous effects. To calculate the fitness effect from the deviation of the phenotype (P) to the environmental optimum (opt) as w = (P −opt)^2^ and the additive genetic variation as V_A_ = Σ2p_i_q_i_z_i_^2^.We ran 100 replicates per scenario and counted the proportion of replicates that went extinct to obtain the extinction rate per scenario.

### Genomic Green Status simulations

We used simulated data from (Jackson *et al*., 2022). Briefly, simulations were performed in SLiM3 (Haller and Messer, 2019) with a non-Wright-Fisher implementation, which considers overlapping generations, age-structure, and customizable offspring generation and migration patterns. During the simulation each time step consists of three stages: reproduction, dispersal (between captive and wild populations, if any), and mortality. Absolute fitness (i.e., probability of survival) was regulated by the carrying capacity and the known aged-based probability of mortality for pink pigeons. The model simulated an exome of 4000 genes of 3.4Kb each with a recombination rate r=1e-4 (no recombination within genes), and a per base mutation rate m=7.5e-8. We modeled neutral genetic variation and a genetic load of ∼15 LEs as observed in the empirical data (see Jackson *et al*., 2022).

We simulated a demographic trajectory that capture the trend observed in the pink pigeon by controlling an overall carrying capacity informed by the inferred (pre-1980s) population size and recorded census trajectories since 1980. We modelled a single panmictic wild population and a captive population founded by 12 individuals (growing to an average of 120 individuals) used to simulate genetic rescue. Four conservation scenarios were simulated across 40 replicate runs each: (1) Counterfactual (no recovery, no genetic supplementation); (2) Demographic rescue (population rebound without genetic rescue); (3) Genetic rescue (genetic supplementation from captive population without demographic increase); and (4) Demographic + genetic rescue, reflecting the actual conservation history in the species. Each replicate tracked nucleotide diversity (π), realized load (sum of homozygous deleterious mutations weighted by s), and extinction over time.

Green Status metrics were calculated for both realized load and nucleotide diversity across the four simulated scenarios. The present-day Green Score (Species Recovery Score, SRS) was compared to hypothetical counterfactuals to calculate four conservation impact metrics: Conservation Legacy (L), Conservation Dependence (D), Conservation Gain (G), and Recovery Potential (R), following the IUCN Green Status framework. Metrics were derived by calculating proportional differences in Green Scores at different timepoints (ancestral, current, 10-year, and 100-year future) under contrasting conservation scenarios.

The Green Score of genetic diversity at time t is expressed relative to ancestral variation and calculated as π_t_ ⁄ π_ancestral_ × 100%. Where π_t_ is the mean nucleotide diversity calculated over the entire population at time t, and π_ancestral_ the mean ancestral nucleotide diversity in the population. The Green Score of realized load is expressed as the realized load in the ancestral variation relative to the realized load in the population at time t. It is calculated for the proportion of runs that survived at time t (P_survived at time t_), giving extinct runs a score of zero as ({2×RL_ancestral_} ⁄ {RL_t_ + RL_ancestral_}) × P_survived at time t_ × 100%. Where RL_ancestral_ and RL_t_ are the mean realized load (RL) in the ancestral population and the population at time t, respectively. In contrast to genetic diversity, the RL_t_ is expected to increase during population size decline because inbreeding and drift expose recessive deleterious mutations in homozygote genotypes. Therefore, to express the Green Score of realized load in negative direction, in this equation the nominator and numerator are switched relative to the Green Score of genetic diversity.

## Data availability

The code used to support the arguments in this perspective are openly available in https://github.com/hmoral/genomic_erosion_perspective.

## Contributions

CvO and HEM conceived the idea. CvO, HEM, CB, LHU, JJG and GS developed the content and idea. HEM, LH and TB performed simulations. CvO and HEM wrote the manuscript with input from all co-authors.

## Acknowledgments

This work was supported by the European Research Council (ERODE: 101078303) and a Royal Society International Collaboration Award, Grant Number: ICA\R1\201194. CvO was funded by the Earth and Life Systems Alliance (ELSA), Norwich Research Park, UK, C.B. is funded by the Wellcome grant WT207492. SAS was funded by NERC ARIES PhD studentship (T209447) at the UEA and a Research Training Support Grant (RTSG; 100162318RA1), TB was funded by a BBSRC (BB/M011216/1) PhD studentship at the UEA. J.J.G. was supported by Research England’s Expanding Excellence in England (E3) Fund, UK Research and Innovation.

## References

Akçakaya HR (2018). Quantifying species recovery and conservation success to develop an IUCN Green List of Species. Conserv Biol 32: 1128–1138.

Al Hikmani H, van Oosterhout C, Birley T, Labisko J, Jackson HA, Spalton A, et al. (2024). Can genetic rescue help save Arabia’s last big cat? Evolutionary Applications 17: e13701.

Araki H, Cooper B, Blouin MS (2007). Genetic effects of captive breeding cause a rapid, cumulative fitness decline in the wild. Science 318: 100–103.

Barton N, Turelli M (2004). Effects of genetic drift on variance components under a general model of epistasis. Evolution 58: 2111–2132.

Beichman AC (2023). Genomic analyses reveal range-wide devastation of sea otter populations. Mol Ecol 32: 281–298.

Bergeron LA, Besenbacher S, Zheng J, Li P, Bertelsen MF, Quintard B, et al. (2023). Evolution of the germline mutation rate across vertebrates. Nature 615: 285–291.

Bertorelle G (2022). Genetic load: genomic estimates and applications in non-model animals. Nat Rev Genet 23: 492–503.

Bolam FC (2021). How many bird and mammal extinctions has recent conservation action prevented? Conserv Lett 14, e12762.

Bonnet T (2022). Genetic variance in fitness indicates rapid contemporary adaptive evolution in wild animals. Science 376: 1012–1016.

Bortoluzzi C, Restoux G, Rouger R, Desnoues B, Petitjean F, Bosse M, et al. (2024). Trends in genome diversity of small populations under a conservation program: a case study of two French chicken breeds. Peer Community Journal 4.

Bortoluzzi C, Wright CJ, Lee S, Cousins T, Genez TA, Thybert D, et al. (2023). Lepidoptera genomics based on 88 chromosomal reference sequences informs population genetic parameters for conservation. bioRxiv: 2023–04.

Brauer CJ, Beheregaray LB (2020). Recent and rapid anthropogenic habitat fragmentation increases extinction risk for freshwater biodiversity. Evol Appl 13: 2857–2869.

Brüniche-Olsen A, Kellner KF, Belant JL, DeWoody JA (2021). Life-history traits and habitat availability shape genomic diversity in birds: implications for conservation. Proceedings of the Royal Society B 288: 20211441.

Buzan E, Guttry C de, Bortoluzzi C, Street N, Lucek K, Rosling A, et al. (2024). Harmonising genomics research excellence and stakeholder needs in conservation management.

Canteri E (2021). IUCN Red List protects avian genetic diversity. Ecography 44: 1808–1811.

Cavill EL, Morales HE, Sun X, Westbury MV, van Oosterhout C, Accouche W, et al. (2024). When birds of a feather flock together: Severe genomic erosion and the implications for genetic rescue in an endangered island passerine. Evol Appl 17: e13739.

Charlesworth B (2009). Fundamental concepts in genetics: effective population size and patterns of molecular evolution and variation. Nat Rev Genet 10: 195–205.

Charlesworth B (2013a). Stabilizing selection, purifying selection, and mutational bias in finite populations. Genetics 194: 955–971.

Charlesworth B (2013b). Why we are not dead one hundred times over. Evolution 67: 3354–3361.

Cook CN, Sgrò CM (2019). Poor understanding of evolutionary theory is a barrier to effective conservation management. Conservation Letters 12: e12619.

Couvet D (2002). Deleterious effects of restricted gene flow in fragmented populations. Conserv Biol 16: 369–376.

Dalton DL, Vermaak E, Smit-Robinson HA, Kotze A (2016). Lack of diversity at innate immunity Toll-like receptor genes in the Critically Endangered White-winged Flufftail (Sarothrura ayresi). Scientific Reports 6: 36757.

Díez-del-Molino D, Sánchez-Barreiro F, Barnes I, Gilbert MTP, Dalén L (2017). Quantifying Temporal Genomic Erosion in Endangered Species. Trends in Ecology & Evolution 33: 176–185.

Dussex N (2021). Population genomics of the critically endangered kākāpō. Cell Genomics 1, 100002.

Dussex N, Kurland S, Olsen R-A, Spong G, Ericsson G, Ekblom R, et al. (2023). Range-wide and temporal genomic analyses reveal the consequences of near-extinction in Swedish moose. Communications biology 6: 1035.

Dussex N, Morales HE, Grossen C, Dalén L, Oosterhout C van (2023). Purging and accumulation of genetic load in conservation. Trends in Ecology & Evolution 38: 961–969.

Exposito-Alonso M, Booker TR, Czech L, Gillespie L, Hateley S, Kyriazis CC, et al. (2022). Genetic diversity loss in the Anthropocene. Science 377: 1431–1435.

Fagan WF, Holmes E (2006). Quantifying the extinction vortex. Ecology letters 9: 51–60.

Falconer DS, Mackay TFC (1996). Introduction to quantitative genetics. Longman: Essex.

Fedorca A, Mergeay J, Akinyele AO, Albayrak T, Biebach I, Brambilla A, et al. (2024). Dealing with the complexity of effective population size in conservation practice.

Femerling G, Van Oosterhout C, Feng S, Bristol RM, Zhang G, Groombridge J, et al. (2023). Genetic load and adaptive potential of a recovered avian species that narrowly avoided extinction. Mo Ecol Evol 40: msad256.

Fogell DJ (2021). Evolution of Beak and Feather Disease Virus across three decades of conservation intervention for population recovery of the Mauritius parakeet. Diversity 13: 584.

Fontsere C, Speak SA, Caven AJ, Rodriguez JA, Wang X, Pacheco C, et al. (2024). Persistent genomic erosion in whooping cranes despite demographic recovery. bioRxiv: 2024–12.

Forester BR, Beever EA, Darst C, Szymanski J, Funk WC (2022). Linking evolutionary potential to extinction risk: applications and future directions. Frontiers in Ecology and the Environment 20: 507–515.

Frankham R (2005). Genetics and extinction. Biological Conservation 126: 131–140.

Frankham R (2008). Genetic adaptation to captivity in species conservation programs. Mol Ecol 17: 325– 333.

Frankham R (2021). Suggested improvements to proposed genetic indicator for CBD. Conserv Genet 22: 531–532.

Frankham R, Ballou JD, Ralls K, Eldridge M, Dudash MR, Fenster CB, et al. (2019). A practical guide for genetic management of fragmented animal and plant populations. Oxford University Press.

Frazer J, Notin P, Dias M, Gomez A, Min JK, Brock K, et al. (2021). Disease variant prediction with deep generative models of evolutionary data. Nature 599: 91–95.

Funk WC, Forester BR, Converse SJ, Darst C, Morey S (2019). Improving conservation policy with genomics: a guide to integrating adaptive potential into US Endangered Species Act decisions for conservation practitioners and geneticists. Conservation Genetics 20: 115–134.

García-Dorado A (2012). Understanding and predicting the fitness decline of shrunk populations: inbreeding, purging, mutation, and standard selection. Genetics 190: 1461–1476.

Gargiulo R, Budde KB, Heuertz M (2024). Mind the lag: understanding genetic extinction debt for conservation. Trends in Ecology & Evolution 0.

Geue JC, Bertola L, Paulette B, Brüniche-Olsen A, da Silva J, DeWoody JA, et al. (2025). Practical genetic diversity protection: an accessible framework for IUCN subpopulation and Evolutionarily Significant Unit identification.

Gilroy DL, Phillips KP, Richardson DS, van Oosterhout C (2017). Toll-like receptor variation in the bottlenecked population of the Seychelles warbler: computer simulations see the ‘ghost of selection past’ and quantify the ‘drift debt’. Journal of Evolutionary Biology 30: 1276–1287.

Goodnight JC (1988). Epistasis and the effect of founder events on the additive genetic variance. Evolution 42: 441–454.

Grace MK, Akçakaya HR, Bennett EL, Brooks TM, Heath A, Hedges S, et al. (2021). Testing a global standard for quantifying species recovery and assessing conservation impact. Conservation Biology.

Grossen C, Guillaume F, Keller LF, Croll D (2020). Purging of highly deleterious mutations through severe bottlenecks in Alpine ibex. Nature communications 11: 1–12.

Grueber CE, Wallis GP, King TM, Jamieson IG (2012). Variation at innate immunity Toll-like receptor genes in a bottlenecked population of a New Zealand robin.

Guillaume F, Rougemont J (2006). Nemo: an evolutionary and population genetics programming framework. Bioinformatics 22: 2556–2557.

Haller BC, Messer PW (2019). SLiM 3: forward genetic simulations beyond the Wright–Fisher model. Mo Ecol Evol 36: 632–637.

Haller BC, Messer PW (2023). SLiM 4: multispecies eco-evolutionary modeling. The American naturalist 201: E127–E139.

Hansson B, Morales HE, Oosterhout C (2021). Comment on ‘Individual heterozygosity predicts translocation success in threatened desert tortoises’. Science 372.

Harrisson KA, Magrath MJ, Yen JD, Pavlova A, Murray N, Quin B, et al. (2019). Lifetime fitness costs of inbreeding and being inbred in a critically endangered bird. Current Biology 29: 2711–2717.

Hedrick PW, Garcia-Dorado A (2016). Understanding inbreeding depression, purging, and genetic rescue. Trends in ecology & evolution 31: 940–952.

Hewett AM, Johnston SE, Morris A, Morris S, Pemberton JM (2024). Genetic architecture of inbreeding depression may explain its persistence in a population of wild red deer. Molecular Ecology 33: e17335.

Hoban S (2023). Genetic diversity goals and targets have improved, but remain insufficient for clear implementation of the post-2020 global biodiversity framework. Conserv Genet.

Hoffmann M (2010). The impact of conservation on the status of the world’s vertebrates. Science 330: 1503–1509.

Hoffmann A (2015). A framework for incorporating evolutionary genomics into biodiversity conservation and management. Climate Change Responses 2: 1–24.

Hohenlohe PA, Funk WC, Rajora OP (2021). Population genomics for wildlife conservation and management. Molecular Ecology 30: 62–82.

Humble E (2022). Conservation management strategy impacts inbreeding and genetic load in scimitar-horned oryx.

IUCN (2004). The IUCN red list of threatened species. Di sponí vel em:< http://www iucn red list org/info/cat e go ries_cri te ria 2001 html> *Aces so em* 12.

Jackson HA, Percival-Alwyn L, Ryan C, Albeshr MF, Venturi L, Morales HE, et al. (2022). Genomic erosion in a demographically recovered bird species during conservation rescue. Conservation Biology 36: e13918.

Jeon JY, Black AN, Heenkenda EJ, Mularo AJ, Lamka GF, Janjua S, et al. (2024). Genomic diversity as a key conservation criterion: proof-of-concept from mammalian whole-genome resequencing data. Evolutionary Applications 17: e70000.

Kardos M (2021). The crucial role of genome-wide genetic variation in conservation. Proc Natl Acad Sci USA 118, e2104642118.

Kardos M, Armstrong EE, Fitzpatrick SW, Hauser S, Hedrick PW, Miller JM, et al. (2021). The crucial role of genome-wide genetic variation in conservation. Proc Natl Acad Sci USA 118: e2104642118.

Kardos M, Zhang Y, Parsons KM, A Y, Kang H, Xu X, et al. (2023). Inbreeding depression explains killer whale population dynamics. Nature Ecology & Evolution 7: 675–686.

Kawakami T, Smeds L, Backström N, Husby A, Qvarnström A, Mugal CF, et al. (2014). A high-density linkage map enables a second-generation collared flycatcher genome assembly and reveals the patterns of avian recombination rate variation and chromosomal evolution. Molecular Ecology 23: 4035–4058.

Keith N (2021). Genome-wide analysis of cadmium-induced, germline mutations in a long-term Daphnia pulex mutation-accumulation experiment. Environ Health Perspect 129: 107003.

Kircher M (2014). A general framework for estimating the relative pathogenicity of human genetic variants. Nat Genet 46: 310–315.

Kleinman-Ruiz D (2022). Purging of deleterious burden in the endangered Iberian lynx. Proc Natl Acad Sci USA 119, e2110614119.

Kyriazis CC (2023). Genomic underpinnings of population persistence in Isle Royale moose. Mol Biol Evol.

Kyriazis CC, Robinson JA, Lohmueller KE (2023). Using computational simulations to model deleterious variation and genetic load in natural populations.

Kyriazis CC, Wayne RK, Lohmueller KE (2021). Strongly deleterious mutations are a primary determinant of extinction risk due to inbreeding depression. Evolution Letters 5: 33–47.

Lacy RC (2019). Lessons from 30 years of population viability analysis of wildlife populations. Zoo biology 38: 67–77.

Laikre L (2010). Genetic diversity is overlooked in international conservation policy implementation. Conservation Genetics 11: 349–354.

Laikre L (2021). Authors’ Reply to Letter to the Editor: Continued improvement to genetic diversity indicator for CBD. Conserv Genet 22: 533–536.

Lighten J, Papadopulos AS, Mohammed RS, Ward BJ, G. Paterson I, Baillie L, et al. (2017). Evolutionary genetics of immunological supertypes reveals two faces of the Red Queen. Nature communications 8: 1294.

Liu X, Milesi E, Fontsere C, Owens HL, Heinsohn R, Gilbert MTP, et al. (2025). Time-lagged genomic erosion and future environmental risks in a bird on the brink of extinction. Proceedings of the Royal Society B: Biological Sciences 292: 20242480.

Lowe WH, Kovach RP, Allendorf FW (2017). Population Genetics and Demography Unite Ecology and Evolution. Trends Ecol Evol 32: 141–152.

Luque GM, Vayssade C, Facon B, Guillemaud T, Courchamp F, Fauvergue X (2016). The genetic Allee effect: a unified framework for the genetics and demography of small populations. Ecosphere 7: e01413.

Lynch M, Conery J, Bürger R (1995). Mutational meltdowns in sexual populations. Evolution 49: 1067– 1080.

Magliolo M (2022). Simulated genetic efficacy of metapopulation management and conservation value of captive reintroductions in a rapidly declining felid. Anim Conserv.

Mathur S, Tomeček JM, Tarango-Arámbula LA, Perez RM, DeWoody JA (2023). An evolutionary perspective on genetic load in small, isolated populations as informed by whole genome resequencing and forward-time simulations. Evolution 77: 690–704.

Matz MV, Treml EA, Aglyamova GV, Bay LK (2018). Potential and limits for rapid genetic adaptation to warming in a Great Barrier Reef coral. PLoS genetics 14: e1007220.

McDonald-Madden E, Baxter PW, Possingham HP (2008). Making robust decisions for conservation with restricted money and knowledge. J Appl Ecol 45: 1630–1638.

McLaughlin CM, Hinshaw C, Sandoval-Arango S, Zavala-Paez M, Hamilton JA (2025). Redlisting genetics: towards inclusion of genetic data in IUCN Red List assessments. Conservation Genetics: 1–11.

Mooney HA, Cleland EE (2001). The evolutionary impact of invasive species. Proc Natl Acad Sci USA 98: 5446–5451.

Mooney A, Ryder OA, Houck ML, Staerk J, Conde DA, Buckley YM (2023). Maximizing the potential for living cell banks to contribute to global conservation priorities. Zoo Biology 42: 697–708.

Morales HE, Groombridge JJ, Tollington S, Henshaw S, Tatayah V, Ruhomaun K, et al. (2024). The genome sequence of the Mauritius parakeet, Alexandrinus eques (formerly Psittacula eques) (A.Newton & E. Newton, 1876). Wellcome Open Res 9: 378.

Morales HE, Norris K, Henshaw S, Tatayah V, Ruhomaun K, Van Oosterhout C, et al. (2024). The genome sequence of the Mauritius kestrel, Falco punctatus (Temminck, 1821). Wellcome Open Res 9: 312.

Morales HE, Van Oosterhout C, Whitford H, Tatayah V, Ruhomaun K, Groombridge JJ, et al. (2024). The genome sequence of the Pink Pigeon, Nesoenas mayeri (Prévost, 1843). Wellcome Open Res 9: 336.

Moran BM (2021). The genomic consequences of hybridization. eLife 10, e69016.

Morris KM, Wright B, Grueber CE, Hogg C, Belov K (2015). Lack of genetic diversity across diverse immune genes in an endangered mammal, the Tasmanian devil (S arcophilus harrisii). Molecular Ecology 24: 3860–3872.

Nadachowska-Brzyska K, Konczal M, Babik W (2022). Navigating the temporal continuum of effective population size. Methods Ecol Evol 13: 22–41.

Nigenda-Morales SF, Lin M, Nuñez-Valencia PG, Kyriazis CC, Beichman AC, Robinson JA, et al. (2023). The genomic footprint of whaling and isolation in fin whale populations. Nature Communications 14: 5465.

van Oosterhout C (2009). A new theory of MHC evolution: beyond selection on the immune genes. Proceedings of the Royal Society B: Biological Sciences 276: 657–665.

van Oosterhout C (2024). AI-informed conservation genomics. Heredity 132: 1–4.

van Oosterhout C, Supple MA, Morales HE, Birley T, Tatayah V, Jones CG, et al. (2025). Genome engineering in biodiversity conservation and restoration. Nature Reviews Biodiversity: 1–13.

Peñalba JV, Wolf JBW (2020). From molecules to populations: appreciating and estimating recombination rate variation. Nat Rev Genet 21: 476–492.

Pérez-Espona S, CryoArks Consortium (2021). Conservation-focused biobanks: A valuable resource for wildlife DNA forensics. Forensic Science International: Animals and Environments 1: 100017.

Pinto AV, Hansson B, Patramanis I, Morales HE, van Oosterhout C (2024). The impact of habitat loss and population fragmentation on genomic erosion. Conserv Genet 25: 49–57.

Ralls K, Sunnucks P, Lacy RC, Frankham R (2020). Genetic rescue: a critique of the evidence supports maximizing genetic diversity rather than minimizing the introduction of putatively harmful genetic variation. Biological Conservation 251: 108784.

Rhymer JM, Simberloff D (1996). Extinction by hybridization and introgression. Annu Rev Ecol Syst 27: 83–109.

Riaño G (2022). Genomics reveals introgression and purging of deleterious mutations in the Arabian leopard (Panthera pardus nimr.

Robinson JA, Kyriazis CC, Nigenda-Morales SF, Beichman AC, Rojas-Bracho L, Robertson KM, et al. (2022). The critically endangered vaquita is not doomed to extinction by inbreeding depression. Science 376: 635–639.

Robinson J, Kyriazis CC, Yuan SC, Lohmueller KE (2023). Deleterious Variation in Natural Populations and Implications for Conservation Genetics. Annual Review of Animal Biosciences 11: 93–114.

Ryman N, Laikre L, Hössjer O (2019). Do estimates of contemporary effective population size tell us what we want to know? Mol. Ecol 28: 1904–1918.

Schmidt C, Hoban S, Hunter M, Paz-Vinas I, Garroway C (2022). The IUCN Red List is not sufficient to protect genetic diversity.

Schulte-Hostedde AI, Mastromonaco GF (2015). Integrating evolution in the management of captive zoo populations. Evol Appl 8: 413–422.

Schwartz KR, Parsons ECM, Rockwood L, Wood TC (2017). Integrating in-situ and ex-situ data management processes for biodiversity conservation. Frontiers in Ecology and Evolution 5: 120.

Segelbacher G (2022). New developments in the field of genomic technologies and their relevance to conservation management. Conserv Genet 23: 217–242.

Shaw RE, Farquharson KA, Bruford MW, Coates DJ, Elliott CP, Mergeay J, et al. (2025). Global meta-analysis shows action is needed to halt genetic diversity loss. Nature 638: 704–710.

Silver L, Farquharson K, Peel E, Gilbert MTP, Morales HE, Hogg CJ (2025). Temporal loss of genome-wide and immunogenetic diversity in a near-extinct parrot. Molecular Ecology 34: e17746.

Smeds L, Ellegren H (2022). From high masked to high realized genetic load in inbred Scandinavian wolves. Mol Ecol.

Somers CM, Yauk CL, White PA, Parfett CL, Quinn JS (2002). Air pollution induces heritable DNA mutations. Proc Natl Acad Sci USA 99: 15904–15907.

Soulé ME (1985). What is conservation biology? Bioscience 35: 727–734.

Soulé ME (1987). Viable populations for conservation. Cambridge University Press.

Speak SA, Birley T, Bortoluzzi C, Clark MD, Percival-Alwyn L, Morales HE, et al. (2024). Genomics-informed captive breeding can reduce inbreeding depression and the genetic load in zoo populations. Molecular Ecology Resources 24: e13967.

Spielman D, Brook BW, Frankham R (2004). Most species are not driven to extinction before genetic factors impact them. Proceedings of the National Academy of Sciences 101: 15261–15264.

Spurgin LG, Richardson DS (2010). How pathogens drive genetic diversity: MHC, mechanisms and misunderstandings. Proceedings of the Royal Society B: Biological Sciences 277: 979–988.

Stoffel MA, Johnston SE, Pilkington JG, Pemberton JM (2021a). Mutation load decreases with haplotype age in wild Soay sheep. Evol Lett 5: 187–195.

Stoffel M, Johnston S, Pilkington J, Pemberton JM (2021b). Genetic architecture and lifetime dynamics of inbreeding depression in a wild mammal. Nature communications 12: 2972.

Terasaki Hart DE, Bishop AP, Wang IJ (2021). Geonomics: Forward-Time, Spatially Explicit, and Arbitrarily Complex Landscape Genomic Simulations. Mol Biol Evol 38: 4634–4646.

Van Der Valk T, Díez-del-Molino D, Marques-Bonet T, Guschanski K, Dalén L (2019). Historical genomes reveal the genomic consequences of recent population decline in eastern gorillas. Current Biology 29: 165–170. e6.

Villemereuil P (2019). Little Adaptive Potential in a Threatened Passerine Bird. Curr Biol 29: 889–894.

Wang X, Fontsere C, Caballero XA, Nielsen SD, Groombridge J, Hansson B, et al. (2025). Genomic erosion through the lens of comparative genomics. bioRxiv: 2025–03.

Waples RS (2024). The Ne/N ratio in applied conservation. Evolutionary Applications 17: e13695.

Waples RS (2025). The idiot’s guide to effective population size. Molecular Ecology: e17670.

Watson R (2019). Summary for policymakers of the global assessment report on biodiversity and ecosystem services of the Intergovernmental Science-Policy Platform on Biodiversity and Ecosystem Services. IPBES Secretariat.

Wilder AP, Supple MA, Subramanian A, Mudide A, Swofford R, Serres-Armero A, et al. (2023). The contribution of historical processes to contemporary extinction risk in placental mammals. Science 380: eabn5856.

Willi Y (2022). Conservation genetics as a management tool: The five best-supported paradigms to assist the management of threatened species. Proc Natl Acad Sci USA 119, e2105076119.

Willis JH, Orr HA (1993). Increased heritable variation following population bottlenecks: the role of dominance. Evolution 47: 949–957.

Willoughby JR (2015). The reduction of genetic diversity in threatened vertebrates and new recommendations regarding IUCN conservation rankings. Biol Conserv 191: 495–503.

Wilson HB, Joseph LN, Moore AL, Possingham HP (2011). When should we save the most endangered species? Ecol Lett 14: 886–890.

